# The Membranous Labyrinth in vivo from high-resolution Temporal CT data

**DOI:** 10.1101/318030

**Authors:** Hisaya Tanioka

**Affiliations:** Department of Radiology, Tanioka Clinic

**Keywords:** 3D imaging, Volume CT, Otologic Diagnostic Technique, Membranous labyrinth, Ear

## Abstract

A prerequisite for the modeling and understanding of the inner ear mechanics needs the accurate created membranous labyrinth. I present a semi-automated methodology for accurate reconstruction of the membranous labyrinth in vivo from high-resolution temporal bone CT data of normal human subjects. I created the new technique which was combined with the segmentation methodology, transparent, thresholding, and opacity curve algorithms. This technique allowed the simultaneous multiple image creating without any overlapping regions in the inner ear has been developed. The reconstructed 3D images improved the membranous labyrinth geometry to realistically represent physiologic dimensions. These generated membranous structures were in good agreement with the published ones, while this approach was the most realistic in terms of the membranous labyrinth. The precise volume rendering depends on proprietary algorithms so that different results can be obtained, and the images appear qualitatively different. For each anatomical question, a different visualization technique should be used to obtain an optimal result. All scientists can create the membranous labyrinth in vivo in real time like a retinal camera.

**Synopsis:** Fusion image of the membranous and the bony labyrinth. Interactive thresholding transparent volume rendering technique is the new method based on the established algorithms. This technique can create the membranous labyrinth in vivo from high-resolution temporal CT data from any scientist.

- The role in the precise volume rendering depended on proprietary algorithms to make 3D histology of the membranous labyrinth from a conventional temporal CT data.
- An intrinsic light transparency of the tissue specification must be taken into account in imaging processing using a specific curve algorithm.
- The 3D in vivo histology can be readily applied to study disease processes and biology of the inner ear without any harmful procedures.

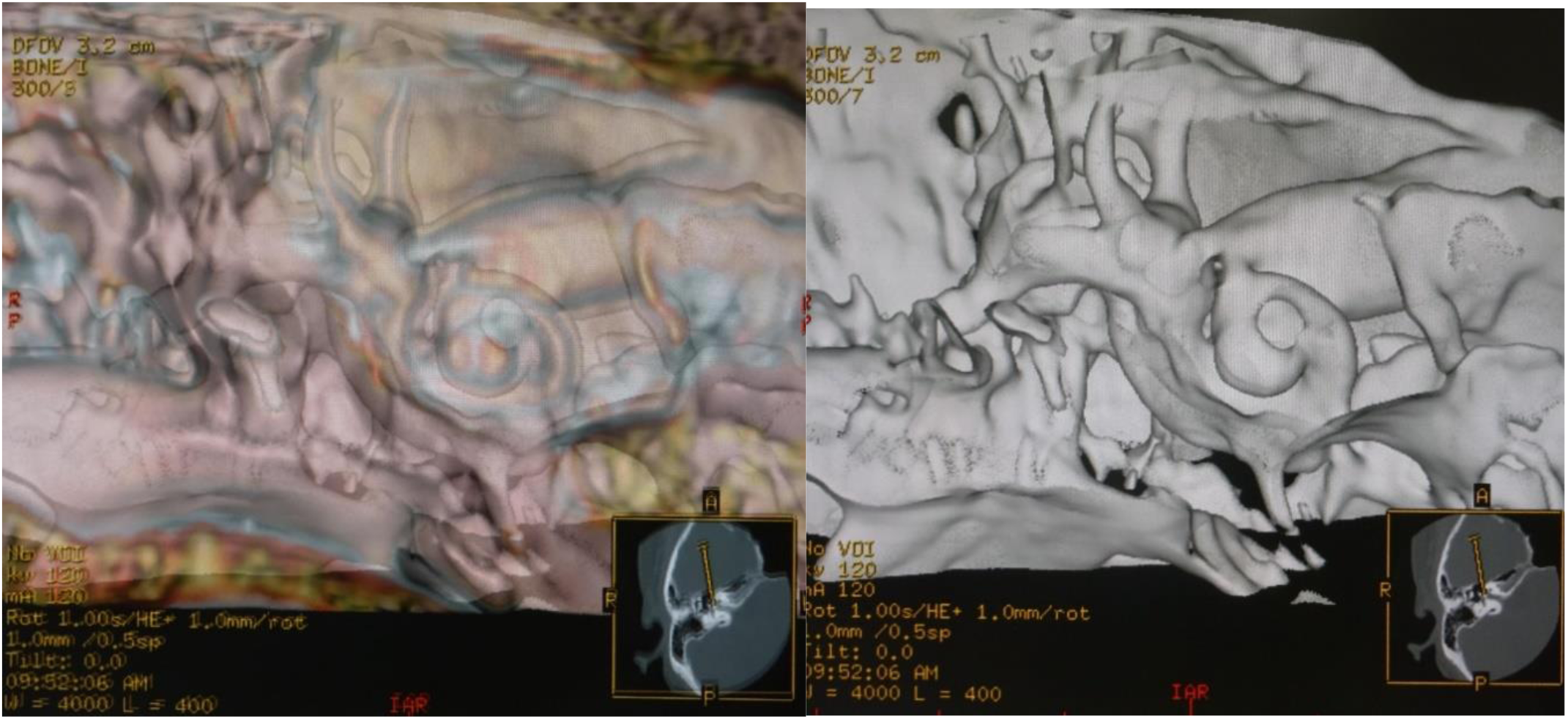

## INTRODUCTION

Developments of radiological modalities and analytic techniques of imaging data have resulted in a steady increase in volume and improvement in the accuracy of the inner ear imaging in the cadaver at a level of a light microscope by using micro-CT and micro-MRI [1]. As scientists have workstations or a PC that downloaded an open-source 3D software, they have come to gain the capabilities for the reconstruction and evaluation of 3D postprocessing from volume data. The 3D reconstructed methods such as maximum intensity projection (MIP), surface rendering, and volume rendering (VR) are the most common computer algorithm used to transform axial volume CT or MRI image data into 3D images. Therefore, the bony labyrinth has been popular to reconstruct from CT or MRI data from the 1990s [1-3]. However, it is impossible to create the membranous labyrinth within the bony labyrinth in vivo. If the soft tissue structures within the osseous labyrinth will be depicted, it will be the image of the membranous labyrinth. In volume rendering (VR) algorithm, high opacity values produce an appearance like surface rendering, which helps display complex 3D relationship clearly. Low opacity value creations can be very useful for seeing a tumor within the lumen [4, 5]. It is possible to create an image of the low density subject behind the high density subject according to the opacity property. Voxel opacity curve algorithms can be adjusted to render tissues within certain density range. Thresholding extracts an interesting region by selecting a range of a target tissue voxel value [6, 7]. Therefore, by combining these algorithms, the low density structure of the membranous labyrinth behind the bone density of the osseous labyrinth can be depicted.

The purpose of this study is to develop and use the volume rendering (VR) algorithms allowing in vivo transparent visualization of the membranous labyrinth from high-resolution CT data sets of the human temporal bone.

## MATERIALS AND METHODS

### Subjects

Imaging data from 10 volunteers (5 men and 5 women, mean age 41.7, range of age: 16-62) who visited Tanioka Clinic for their regular checkup and wished to inspect the head and temporal bone regions were obtained. These subjects had no known affliction of temporal bone and no history of hearing problem, or other systems related to auditory and vestibular systems and normal findings on both CT exams and physical exam by their checkup. This study was conducted under the approval of the clinic institutional review board with the 1964 Helsinki Declaration and its later amendments or comparable ethical standards.

### CT Protocol

All examinations were performed with the spiral CT scanner (ProSpeed AI; General Electric Systems, Milwaukee, Wis., USA) by using an axial technique with 120 kV, 60 −120 mA, and 60 seconds scanning time. The section thickness was 1mm and an interval of 0.5 mm overlapping. A pitch was 1.0. The axial images were reconstructed with a high-resolution bone algorithm in steps of 0.5 mm, and a field of view (FOV) of 96 × 96 mm by using a 512X512 matrix. The range of dose length projection (DLP) was 40 mGy/cm to 55 mGy/cm.

### Volume Rendering (VR)

Most VR technique will be divided into two types [4-7]. One is thresholding, or surface-based methods. The other is percentage- or transparent volume based techniques. Certain properties (opacity, color, shading, brightness) are assigned to each voxel value. The most important property is opacity or attenuation, which enables the user to hide certain objects or make them transparent. Thresholding extracts a region of interest by selecting a range of voxel values that presents a specific tissue or anatomical feature. Voxel opacity curve algorithms can be adjusted to render tissues within certain density range. In this study, the algorithms were made from a thresholding method with a transparent volume based technology under voxel opacity curve algorithms. The multitude of user-definable parameter settings will preserve the accuracy of VR for the evaluation of the membranous labyrinth, as the selection of window width (WW), window level (WL), brightness, and opacity differ for each observer. Hence, it is necessary to set each parameter, and to choose which type of a voxel opacity curve algorithm beforehand for 3D image creation.

### Parameter Setting

#### Threshold Value

All measurements were carried out using the CT workstation (Advantage version 2.0; GE Medical Systems, Milwaukee, Wis). To define the optimal threshold CT values to create 3D images of the membranous labyrinth, CT values within the edge of the cochlea and vestibule, and the modioulus were measured using 0.5-mm-thick axial HR-CT image slices at the level through the oval window, the cochlea, and the vestibule. The imaging condition was as follows; WW was 2000HU, and WL was 200HU. Pixel-level analysis of the DICOM viewer yield actual CT values of them. The results were shown in Table 1.

**Table 1.**
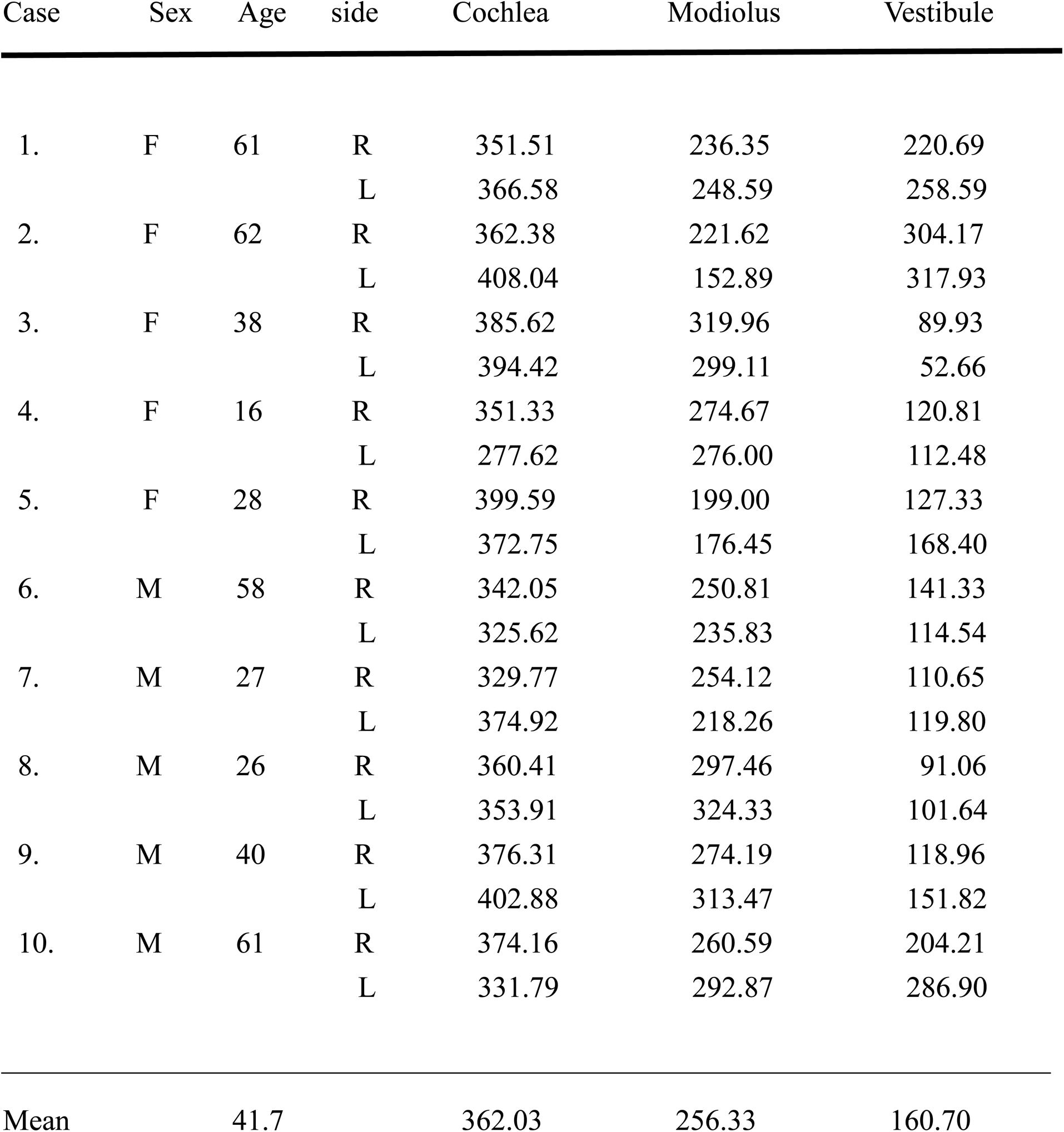
X-ray absorption values of the inner ear at the level of 1^st^ to 2^nd^ turn of the cochlea

#### Statistical Analysis

Normality of distribution was tested using the Graphical methods. Goodness of fit test for normality was evaluated by Chi-square statistics [8]. The results were shown in table 2.

**Table 2:** The results of goodness of fit test for normal distribution

|  | Cochlea | Modiolus | Vestibule |
| --- | --- | --- | --- |
| rank | 6 | 6 | 6 |
| mean | 362.033 | 256.328 | 160.695 |
| SD | 31.301 | 46.899 | 77.771 |
| variance | 1008.819 | 2286.363 | 6044.256 |
| degree of freedom | 3 | 3 | 3 |
| $\chi^2$ | 0.66275 | 1.97023 | 4.36420 |
| significance | NS | NS | NS |
| precise P value | 0.88193 | 0.57866 | 0.22472 |

#### Define Threshold Value

The statistical analysis revealed that the results were suitable for a normal distribution. The results were given as mean±SE. The cochlea was 362.1 ± 13.7 HU, the modiolus was 256.3 ± 20.6 HU, and the vestibule was 160.7 ± 34.1 HU. The results were presented as median values and 95% confidential intervals (CI). These results suggested that the optimal CT values of the cochlea were about through 348 to 376 HU, of the modiolus were about through 236 to 277 HU, and of the vestibule were about through 124 to 197 HU. The cochlear duct CT value was the soft tissues within the fluid components in the cochlea. By subtracting the modiolus value from the cochlear value, the cochlear duct value was obtained. The calculated value was in the range of about 71 to 140 HU, and the average was 105 HU. To know the state of the modiolus together, the highest value of the modulus was taken as the high threshold value. From the anatomical components, the medullary sheath contained fat. That is, the low absorption value was a fat value of −100 HU, and the high absorption value was 280 HU.

#### Opacity Value

Opacity values are determined to define the relative transparency of each material. High opacity values produce an appearance like surface rendering, which helps display complex 3D relationship clearly. Low opacity value creations can be very useful for seeing a subject within the lumen [4-7]. Hence, it is possible to observe the low density (high opacity) subject behind the high density (low opacity) subject according to the property of opacity. As for as the cochlea is concerned, by assigning a low density (high opacity) to the soft tissue and a high density (low opacity) to the modiolus, the soft tissue becomes clearly visible among its semi-transparent surroundings. Therefore, the opacity value is defined by the ratio of the CT values between the two subjects.

That is, the following equation holds.

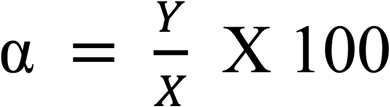

where α is the opacity value, Y is the low CT density value, and X is the high CT density value.

The range of the opacity value was 25.6 to 59.3. The average value was 41.0. The opacity value was set to 40%.

#### Brightness Value

The brightness value was defined as constant value of 100%.

### Algorithm Procedure

#### Select Opacity Curve Algorithm

A voxel opacity curve algorithm can be adjusted to render tissues within certain density range. There are three opacity curve algorithms to reconstruct 3D images of the membranous labyrinth as follows.

1. Upward opacity curve algorithm As the structures over the high threshold value will show a constant high opacity value, under the lower threshold value is 0% opacity value. Thus, it may be used to display bright structures such as bone or vessels in CT data sets [6]. That is, this opacity curve is suitable to delineate soft tissue structures on the bony labyrinth like Fig. 1.
2. Trapezoid opacity curve algorithm This opacity curve is used to display structures with voxel values within the defined range [7]. This is suitable for the surface structure of the membranous labyrinth like Fig. 2, 3, and 4.
3. Downward opacity curve algorithm In case of this opacity curve type, it is opposite to an upward opacity curve type. This opacity curve type can be used to display dark structures such as airways, or vessels in black blood MRA technique [9]. It can also be used with a cut plane to display the lumen of a vessel as etched in the plane. This means that lower X-ray absorption organs are distinguished from higher X-ray absorption subjects. So, this opacity curve is suitable for fibrotic structures in the membranous labyrinth like Fig. 5.

**Fig. 1:**
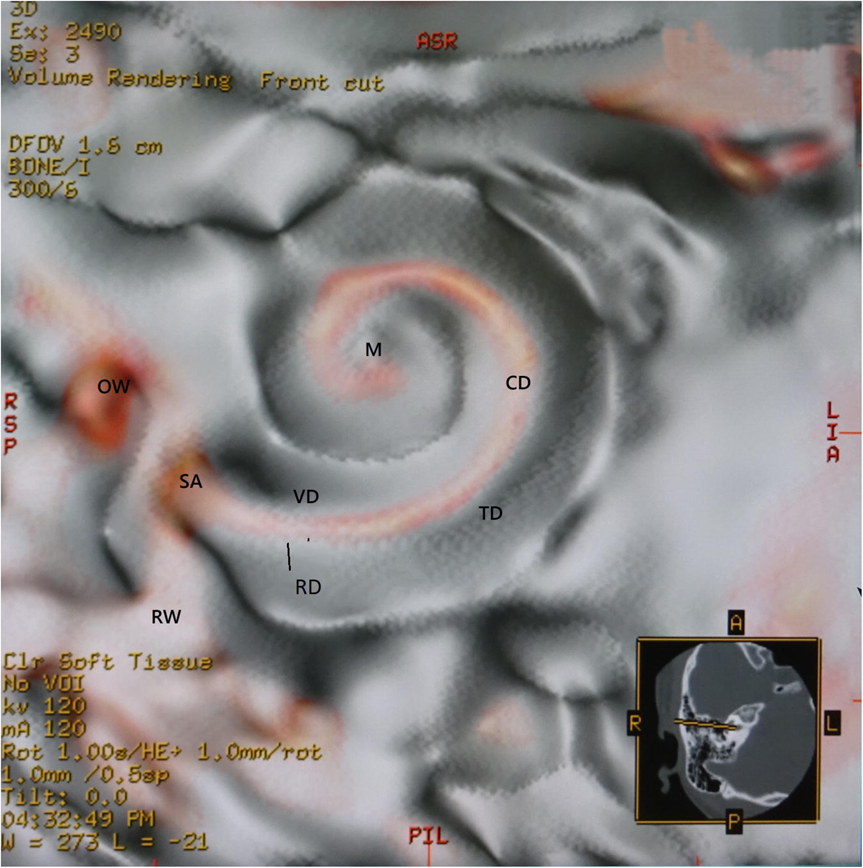
The cross section perpendicular to the modiolus to observe the entire cochlear duct in 16-year-old female. This image is made by an upward curve with a merged colored algorithm, and the opacity value is 40 %. The cochlear soft tissues are a spring duct between the tympanic duct and the vestibular duct. The image is like removing the lid of the cochlea, and looks like the whole organ of Corti with the ossoeus lamina. The DFOV is 16 mm. Abbreviations: CD = the cochlear duct. TD = the tympanic duct. VD = the vestibular duct. RD = the ductus reuniens. M = the modiolus. SA = the saccule. RW = the round window. OW = the oval window.

**Fig. 2:**
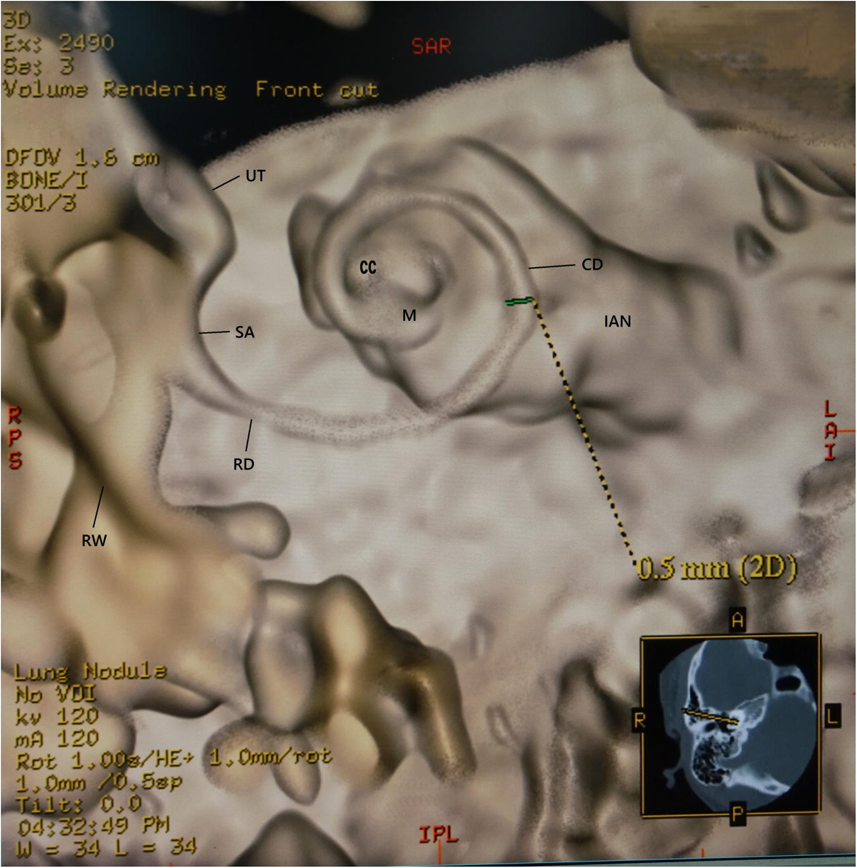
The region of measured diameter at one quarter of the basal turn of the cochlear duct. The same view and subject as in Fig. 1. This image is made from a trapezoid curve with a non-merged colored algorithm, and the opacity value is 40 %. The cochlear duct is winding along the modiolus. The shape of the modiolus shows conical axis. The DFOV is 16 mm. Abbreviations: CD = the cochlear duct. M = the modiolus. RD = the ductus reuniens. IAN = internal auditory nerve. SA = the saccule. UT = the utricle. RW = the round window. CC = the cupular cecum.

**Fig. 3:**
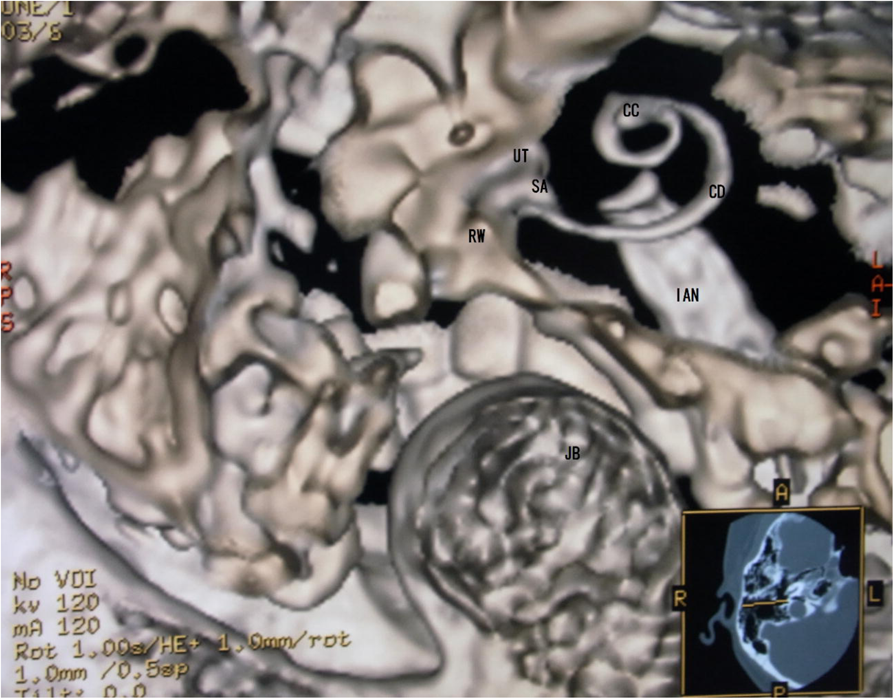
Superiolateral view of the cochlea in 28-year-old female. This image is made by a trapezoid curve with a non-merged colored algorithm and 40 % opacity value. The cochlear duct has a shape of a triangular spiral tube, and the end of the duct showed like a snake head is the cupular cecum. The saccule shows an egg shape from this view. The DFOV is 32 mm. Abbreviations: CD = the cochlear duct. CC = the cupular cecum. SA = the saccule. UT = the utricle. RW = the round window. IAN = internal auditory nerve. JB = the jugular bulb.

**Fig. 4:**
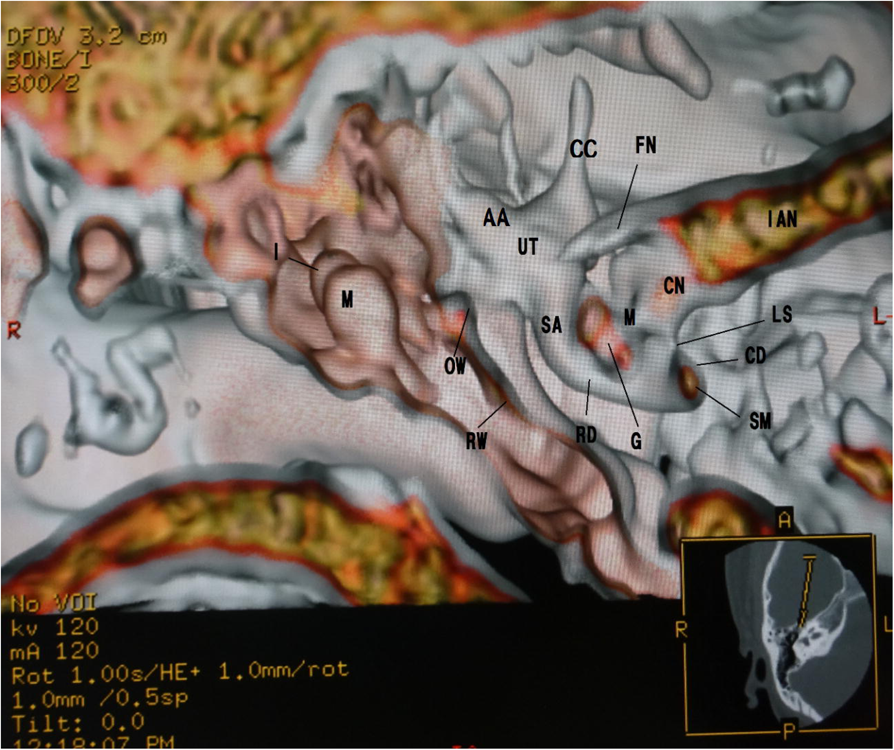
Anterior cochlear cut image in 16-year-old female. This image is made by a trapezoid curve with a merged colored algorithm and 40 % opacity value. The cochlear duct is a spiral duct and a triangular shape in the cross section. This duct is suspended between the osseous spiral lamina of the modiolus. The spiral membrane is located within the cochlear duct. The spiral ganglion is situated along the modiolus. The DFOV is 32 mm. Abbreviations: CD = the cochlear duct. SM = the spiral membrane. LS = the osseous lamina. M = the modiolus. CN = cochlear nerve. G = the spiral ganglion. IAN = internal auditory nerve. FN = facial nerve. SA = the saccule. UT = the utricle. AA = the anterior ampulla. CC = the common crus. RW = the round window. OW = the oval window. M = the malleus. I = the incus.

**Fig. 5:**
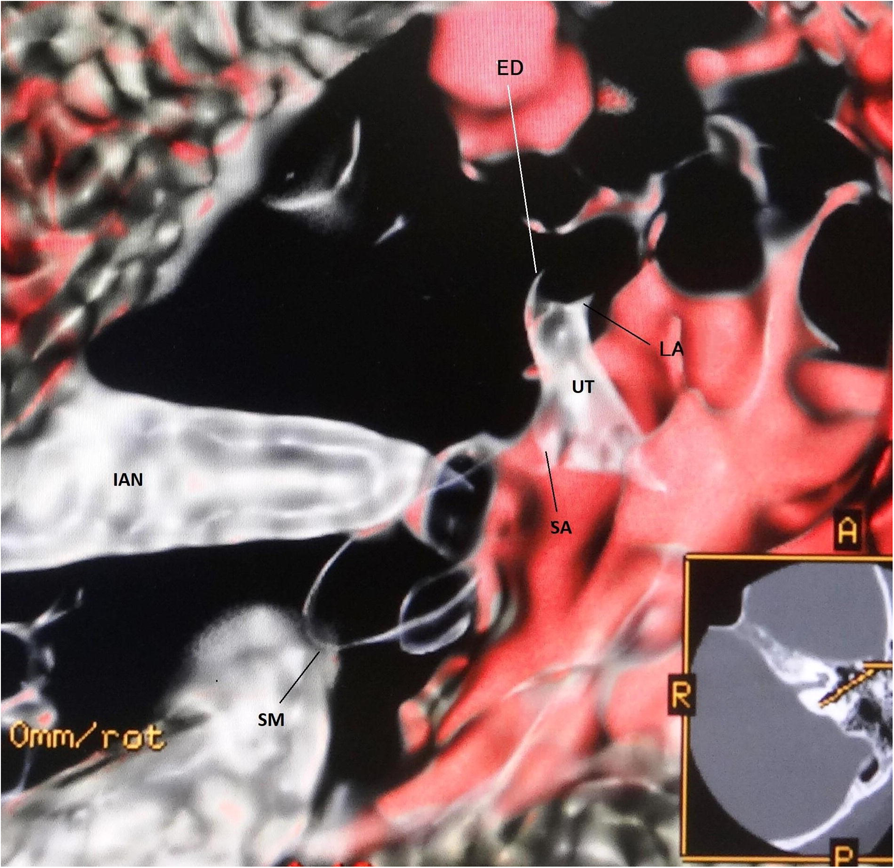
Superior view of the cochlea in 58-year-old male. This image is made by a downward curve with a merged colored algorithm. The opacity value is 40 %, and the high threshold value is 280 HU. The spiral membrane is a spiral winding within the cochlear duct. The diameter of the cochlear duct in Fig. 2 and 3 is wider than that of the spiral membrane as in Fig. 5. The width of a spiral winding becomes wider from the base through the apex. The DFOV is 32 mm. Abbreviations: SM = the spiral membrane. UT = the utricle. SA = the saccle. IAN = internal auditory nerve. ED = the endolymphatic duct. LA = the lateral ampulla

#### Color Shading Mode Algorithm

When the color shading method is used to create images, the image is shaded based on the orientation of the surfaces in the voxels that contribute to given pixel, and colored according to the value of these voxels [4-7]. Color shading is surface shading, and it can be seen in the final relationship that basing the opacity functions in the color space effectively revealed structures within volumes.

#### Applying Colors Algorithm

There are two techniques to create 3D images by applying color methods as follows [4-6].

1. Non-merged 3D image When a non-merged image is colored, everything in it takes on the same color like Fig. 2 and 3. To do this, select the image to be colored, then chose the desired color. Coloring of a non-merged image is typically done in preparation for a 3D merging operation. This makes distinguishing the merge objects easier.
2. Merged 3D image When the coloring is applied to a merged 3D view, each merged 3D model can be colored independently of the others like Fig. 1, 4, and 5.

#### Simple Segmentation Algorithm

As the target organ data existed in the inner ear, roughly simple segmentation of the temporal bone was required using a front cut method.

#### Postprocessing

After transferring the image to the CT workstation (GE Medical Systems, Milwaukee, Wis., USA), 3D visualization based on interactive direct volume rendering was made using GE Advantage Navigator software. The direct volume rendering considers some of the image data, so roughly simple explicit segmentation prior to the visualization process was required using a front cut method. The display field of view (DFOV) was 16-32 X16-32 mm. As a result of this software, both color and opacity values were adjusted interactivity to delineate all structures related to the membranous labyrinth in real time under the threshold value. After an appropriate setting, the front cut for optimal delineation of the objective structures was defined on axial, coronal and sagittal projection, the color and opacity table could be stored and used for further studies. Owing to the simple acceleration of the visualization process, the whole procedure was performed in less than 1 minute. The standardized user ensured intuitive manipulation of any object in real time. The software allowed both distance measurements and volume directly within the 3D scene. This software was almost the same as the open-source OsiriX software [10] (available at http://www.osirix-viewer.com) or LiveVolume (available at http://www.livevolume.com) [11].

#### Quantitative Image Analysis based on the results of previous histological literatures

To ascertain whether the 3D created images were consistent with anatomical findings, the cochlear duct width was measured at the position as shown in Fig. 2, and compared with the results of previous histological literatures. The result of the cochlear duct width was 0.40 to 0.60 mm, and then mean ± SD value was 0.50 ± 0.08 mm. According to the previous literatures, Retzius reported as follows; the height of the lateral wall was from 0.59 to 0.3 mm, the width of the spiral limbus was from 0.22 to 0.23 mm, and the width of the basilar membrane was from 0.21 to 0.36 mm [12]. Wever and Laurence reported that the width of the basilar membrane was between 0.498 to 0.80 mm [13]. That is, the height of the lateral wall was about 0.50mm, and the width of the floor was about between 0.40 to 0.60 mm. The difference between the results of previous literatures and of this study was assessed for significance using a Wilcoxon rank sum test. A p value of < 0.05 was regarded as significant.

## RESULTS

### Volume Rendering Parameters

The threshold value was −100 to 280 HU, the opacity value was 40%, and the brightness value was 100 %.

### Modeling

The different 3D membranous histological images were created by three different voxel opacity curve algorithms with a merged or non-merged colored technique. The quite different 3D images were made from the different opacity curve algorithm, though Fig. 1 and Fig. 2 were created from the same direction of the same subject. These created images were the same that of the published anatomical books and literatures [12-17]. The relation between the created image and the algorithm was as follows.

1. The soft tissues of the membranous labyrinth on the bone were created from an upward opacity curve with a merged color algorithm. (Fig. 1)
2. The surface figure of the membranous labyrinth was made from a trapezoid opacity curve with a non-merged colored algorithm. (Fig. 2 and 3)
3. The transparent membranous labyrinth was created from a trapezoid opacity curve with a merged colored algorithm. (Fig. 4)
4. The fibrotic minute structures in the membranous labyrinth was reconstructed from a downward opacity curve with a merged colored algorithm. (Fig. 5)

### Quantitative Image Analysis

The measured result in the 3D created membranous labyrinth was 0.40 to 0.60 mm. A significant correlation in comparison with the results of the previous literatures was found p = 0.0048.

## DISCUSSION

The aim of this study was to investigate the non-invasive 3D creation method of the membranous labyrinths in vivo. According to this study, the precise volume rendering depends on proprietary algorithms so that different results can be obtained, and the images appear qualitatively different. For each anatomical question of the membranous labyrinth, a different visualization technique should be used to obtain an optimal result.

### On the question of whether it is an anatomically accurate model or not

The measured result of the width of the cochlear duct in this study was nearly the same as the histologically measurement values in the previous literatures [12.13]. This finding supports that these created 3D images will be accepted by morphologists.

### The problem of Threshold value

Defined as the threshold value was obtained from the measured value in this CT exam, this value cannot be instantly adapted to other CT equipment. But this defined threshold value will be referred when setting a threshold value with different CT machines.

### The Problem of Resolution

The resolution of 3D images is primarily determined by the resolution of the original acquisition images and the inter-slice distance. Although the pixel size of the original 2D image is large, the voxel size of the reconstructed display 3D image is smaller in the practical 3D image. This cause will be based on an algorithm of 3D configuration [18]. It is not possible to identify details within the exam with dimensions in the order of or less than the inter-slice distance with any degree of reliability. Therefore, at all times, it remains the responsibility of the scientist to determine whether the inter-slice distance used for a particular exam is acceptable. Despite the intra-slice distance is 0.5 mm in this study, the created 3D histological images were in good agreement with published ones. However, the more the amount of accurate thin section 2D data, the more accurate 3D reconstructed images will be created.

## Conclusions

The new interactive transparent volume rendering algorithm is created in combination with thresholding, opacity curve, and transparent algorithms. This new technique allows the simultaneous multiple image creating without any overlapping regions in the inner ear, and all scientists can use easily. Hence, this technique can study the membranous labyrinth in vivo in real time like a retinal camera.

## Compliance with Ethical Standards

### Guarantor

The scientific guarantor of this publication is Hisaya Tanioka.

### Conflict of Interest

The author of this manuscript declares no relationship with any companies, whose products or services may be related the subject matter of the article.

### Funding

The author states that this work has not received any fundings.

### Statistics and Biometry

No complex statistical methods were necessary for this paper **Informed consent:** Written informed consent was obtained from all individual participants in this study. **Ethical approval:** All procedures performed in studies involving human participants were accorded with the Ethics Committee of Tanioka Clinic, Japan, and with the 1964 Helsinki Declaration and its later amendments or comparable ethical standards. Institutional Review Board approval at Tanioka Clinic was obtained.

### Methodology

Prospective; experimental, performed at one institution.

